# VarSCAT: A computational tool for sequence context annotations of genomic variants

**DOI:** 10.1101/2022.11.11.516085

**Authors:** Ning Wang, Sofia Khan, Laura L. Elo

## Abstract

The sequence contexts of genomic variants play important roles in understanding biological significances of variants and potential sequencing related variant calling issues. However, methods for assessing the diverse sequence contexts of genomic variants such as tandem repeats and unambiguous annotations have been limited. Herein, we describe the Variant Sequence Context Annotation Tool (VarSCAT) for annotating the sequence contexts of genomic variants, including breakpoint ambiguities, flanking sequences, variant nomenclatures, adjacent variants, and tandem repeats with user customizable options. Our analysis demonstrate that VarSCAT is more versatile and customizable than current methods or strategies for annotating variants in short tandem repeat (STR) regions. Variant sequence context annotations of high-confidence human variant sets with VarSCAT revealed that more than 75% of all human individual germline and clinically relevant insertions and deletions (indels) have breakpoint ambiguities. Moreover, we illustrate that more than 80% of human individual germline small variants in STR regions are indels and that the sizes of these indels correlated with STR motif sizes. VarSCAT is available at https://github.com/elolab/VarSCAT.

**Author Summary:** The sequence contexts have significant impacts on the biological and technical aspects of genomic variants. The sequence contexts, such as tandem repeats or nearby indels, may increase the mutation rate of a region compared to other genome regions. Besides, variants in specific sequence contexts like STRs may also have distinguished biological consequences, which can lead to certain human diseases and thus they may be used as biomarkers for disease diagnosis and treatments. Moreover, potential ambiguous variant representations such as equivalent or redundant indels are also related with their sequence contexts, which may complicate variant harmonization from different sources. Our previous study demonstrated that more than half of false positive indel calls detected through next generation sequencing data are related with STRs. Thus, the sequence contexts of genomic variants are important and cannot be ignored. However, the current methods or strategies for assessing the sequence contexts of genomic variants are limited and not feasible to use. Here, we developed a computational tool VarSCAT for sequence contexts annotation of genomic variants. Our tool provides diverse sequence contexts annotations providing users information to further explore the variants of their interests. By applying VarSCAT to high confidence human variant sets, we demonstrate the influence of sequence context of genomic variants and emphasize the importance of sequence context assessment.

## Introduction

Genomic variants can influence the fundamental biological processes. Germline variants, which occur in germ cells and can be transmitted to subsequent generations, are the major source of heritable genetic variation. Somatic variants, which occur in any cells except germ cells, can only be transmitted to their daughter cells [1]. These genomic variants may be resulting in gain or loss of functions of their encoded proteins and cause some diseases such as cancers, or show associations with some certain phenotypes through gene regulation networks [2,3]. Evidence has been shown that the sequence contexts of genomic variants can have biological and technical influences on the properties of variants. Several studies have shown that the mutation rate of variants is affected by nearby nucleotide patterns and genomic features such as GC contents and CpG islands [4–6].

Another study illustrated that the single nucleotide mutation rate increased when nearby insertions and deletions (indels) were present [7]. Short tandem repeats (STRs), which are mainly caused by the DNA strand slippage and that compose approximately 3% of the human genome, are known as important sequence context features of genomic variants [8,9]. In humans, STR regions have relatively high mutation rates compared with single nucleotides, making them among the fastest-evolving DNA sequences [10–12]. The evolutionary mutational pattern of STRs usually increases or decreases by one repeat motif at a time, but the pattern can also be complicated and heterogeneous [13]. Variants in STR regions may also play important roles in the molecular and cellular functions associated with human health and diseases. For examples, the expansion of a CAG trinucleotide repeat in the *HTT* gene can causes Huntington’s disease, and an increased copy number of a CGG trinucleotide repeat in the *FMR1* gene can causes fragile X syndrome [14–16]. Microsatellite instability (MSI) is found in tumor tissues of many cancer types, and it contains STR mutations caused by impaired DNA repair system. MSI can be used as biomarkers for cancer diagnosis and treatment [17–20]. One *in vitro* research demonstrated that the sequence context of frame-shift indels in STR regions may promote their tolerance via bypass of transcriptional or translational errors [21].

Besides, the sequence contexts of genomic variants can also cause difficulties in next generation sequencing (NGS) data analysis, such as the breakpoint ambiguity of indel calling [22,23]. Breakpoint ambiguity is caused by microhomological subsequences surrounding an indel site, which create an equivalent region for the indel, making it impossible to identify the exact breakpoint position of the indel but to locate the breakpoint at a 5’-aligned position [22,23]. Breakpoint ambiguities may cause potential problems for downstream annotations, such as the Human Genome Variation Society (HGVS) nomenclature of variants, which recommends a 3’-aligned position but may lead to redundancies of biologically equivalent indels [24,25]. Study [22] illustrated that indel breakpoint ambiguities can affect the sensitivity of indel calling and suggested the unambiguous annotation of an indel, which should have a single coordinate and an equivalent indel region, depending on its sequence context. Another study [23] showed that 40% of deletions ≥ 32 bp in the human genome cannot be identified with unique positions by alignments of 100 bp sequencing reads. In our previous study [26], we found that more than half of the false positive indels detected by a variety of variant calling methods using NGS data were located in the STR regions. These results highlight that the indel breakpoint ambiguity caused by similar local sequence contexts cannot be ignored. The low complexity and highly similar sequence contexts of STRs may also cause technical problems in NGS. The relatively short reads of the Illumina sequencing platform [27] cannot fully resolve long STR regions because the repeat regions may be longer than the length of the reads [28]. Although the single molecule real-time sequencing platforms, such as Pacific Biosystems [29] and Nanopore sequencing [30], have longer read lengths, which can successfully span large regions of STRs, the relatively high error rates and additional costs limit their further application [28].

A variety of methods and resources can be utilized to assess the sequence contexts of variants in the human genome. The sequence contexts of variants can be assessed by analyzing the reference sequence or calling variants in specific sequence contexts. For example, Benson et al. [31] developed the Tandem Repeats Finder (TRF) programme by applying a probalilistic model to locate and display STRs with DNA sequences in FASTA format. The ‘Simple Repeats’ and the ‘Microsatellite’tracks of the current version of the University of California Santa Cruz’s (UCSC’s) Genome Browser were created using TRF [32]. Krait is a computational tool for detecting different types of STRs with user-defined parameters from a DNA sequence in FASTA format [33]. The ‘TandemRepeat’ function of the Genome Annotation Toolkit (GATK) variant annotation module can annotate copy numbers and repeat motifs from a input Variant Call Format (VCF) file if the variant is located in perfect STRs [34]. UPS-indel is a web service with an additional command-line interface that uses a universal positioning system to mark potential breakpoint ambiguities and determine the unique coordinates of indels from VCF input files [25]. SeqTailor is a web server for extracting DNA or protein sequences with reference and alternative alleles directly from VCF input files, which is useful for retrieving information about the sequence contexts of variant sites [35]. The Variant Tools software has several functions for variant analysis, one of which can output flanking bases of reference and alternative alleles from VCF or custom-format input files [36]. HipSTR [37], STRetch [38], exSTRa [39], and GangSTR [40] are computational tools that take either short-read sequencing FASTQ files or read alignment files as the inputs to analyze tandem repeats in pre-selected STR regions. These pre-seoected STR regions may require additional steps and prior knowledge from users to produce them in specific formats.

However, currently, no tool can comprehensively annotate the sequence contexts of variants, including breakpoint ambiguity, flanking sequences, HGVS nomenclature, and tandem repeats, directly from VCF files in a high-throughput manner. For example, the existing STR annotation methods, using STR locations in BED format to annotate variants in a VCF file, may require several tools or steps; thus, a computational pipeline may be needed to annotate variants with pre-selected STR locations. Also, due to the choice of parameters, these methods may only focus on certain types of STRs, limiting the further analysis, and some tools, such as SeqTailor and Variant Tools, have functions that visualize sequence contexts around variant sites but cannot give information about local sequence complexity. Herein, we present VarSCAT, which is a variant sequence context annotation tool with various functions for studying the sequence contexts around variants and annotating variants with breakpoint ambiguities, flanking sequences, HGVS nomenclature, distances between adjacent variants, and tandem repeat regions.

## Results

### An overview of VarSCAT

We developed a command-line-based computational tool VarSCAT for annotating variant sequence contexts. VarSCAT takes a VCF file and a reference sequence as inputs to provide information about the sequence contexts of variants with single command lines (Fig 1). The variant normalization module is a pre-processing module which first splits any potential multiallelic variants into biallelic variants and then normalizes all input variants parsimonious and left aligned. The ambiguous variant annotation module is able to output breakpoint ambiguity and the affected regions of variants, the HGVS nomenclature of variants based on genomic coordination, distances between adjacent variants, and flanking bases of both the reference and alternative alleles, even those containing ambiguous breakpoint regions. If genomic coordinates are given, VarSCAT can output the subregion of the reference and mutated sequences that contain variants as well as complementary sequence of the mutated sequence. With the tandem repeat region variant annotation module, VarSCAT can annotate the local sequence contexts around variants and annotate putative tandem repeat regions that contain variants with default or user-defined parameters for purity, composition, and size of putative tandem repeats.

**Fig 1.**
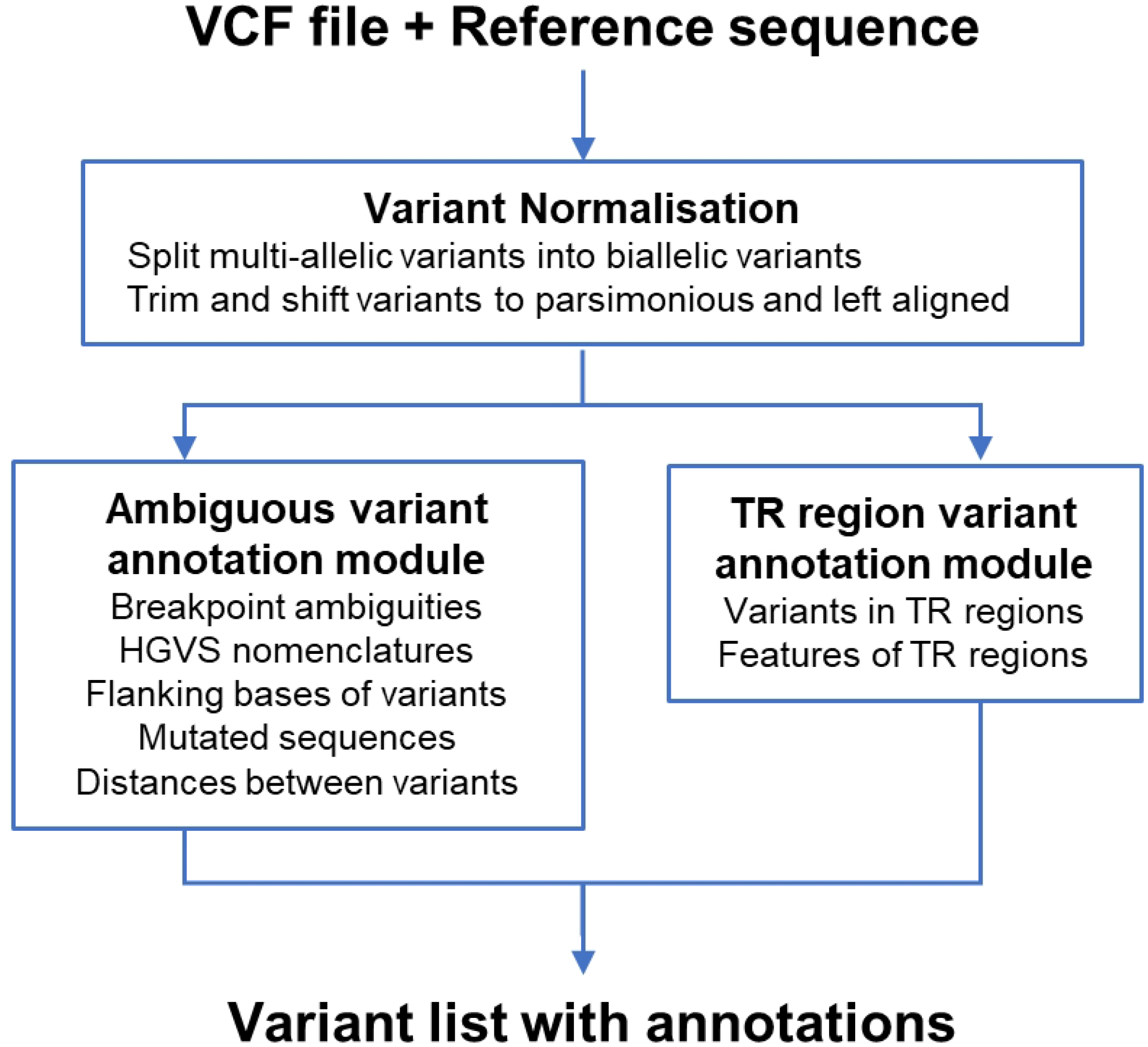
VarSCAT workflow. The input VCF file is first passed to the variant normalization, where the essential information is extracted. The two VarSCAT annotation modules are an ambiguous variant annotation module and a tandem repeat (TR) region variant annotation module, which can be used to annotate the breakpoint ambiguities, flanking sequences, HGVS nomenclature, distances between adjacent variants, and tandem repeat regions of input variants, respectively. The VarSCAT results, generated with a single command line, are written into a text file that lists of the variants with their corresponding annotations.

To assess our newly developed tool, we used variants based on human reference assembly GRCh38 from the ClinVar database [46], Platinum Genome [47], the National Institute of Standards and Technology’s Genome in a Bottle (GIAB) [48,49], and the 1000 Genomes Project [50,51] to study the breakpoint ambiguities of indels and small variants (1–50 bp) in STR regions. Due to the varying algorithmic parameters used in different studies for STR detection, such as the minimum length of an STR and the tolerance of mismatches and indels between STR units, the definitions of an STR may vary widely and lead to highly variable interpretations [52–54]. In our study, we restricted our analysis to perfect STRs with motif sizes of 1–6 bp based the common definitions in the literature [38,55–57]. We set the minimum length of a STR region to 10 bp according to a computational and experimental study about DNA polymerase-mediated strand slippage rates [58], and we set the minimum copy number to 10 for mononucleotide STRs, 5 for dinucleotide STRs, and 4 for tri- to hexanucleotide STRs, according to [58], a meta-analysis [59], and an *in silico* study about microsatellite distributions in the human genome [60] (see S1 file: Sections S1–S3).

### Benchmarking VarSCAT against other methods for annotating variants in STR regions

We analyzed high-confidence small variants located in the human reference genome GRCh38 chromosome 1 of two human individuals (HG002 and HG006) from the GIAB Consortium. The current strategies for annotating variants in STR regions are as follows: 1) directly annotate variants in STR regions with a reference genome, 2) download ready-made STR annotations and then use the annotation tools to annotate variants, or 3) detect STR regions with a reference genome and then use the annotation tools to annotate variants. To incorporate various annotation strategies into benchmarking, we selected GATK (v4.1.9.0, ‘TandemRepeat’ function) [61] for direct annotation, Krait (v1.3.3) [33] for detecting and annotating perfect STRs against a reference genome, and TRF [31] and RepeatMasker [62] from UCSC Genome Browser’s ‘Simple Repeats’ and ‘RepeatMasker’ tracks (download date 11 February 2022) to represent the ready-made STR annotations of a reference genome. For annotations of the reference sequence GRCh38 chromosome 1, we used ANNOVAR (version: ‘$Date: 2019-10-24’) [63] as the annotation tool to annotate variants of GIAB HG002 and HG006 using VCF files. After collecting variants located in STR regions using different annotation methods, we used Venn diagrams to view the overlaps among the different annotation sets (S1 file: Section S2).

The results demonstrated that VarSCAT could annotate a larger collection of variants located in the STR regions than other annotation methods (Fig 2). VarSCAT and GATK ‘TandemRepeat’ shared a large proportion of annotated variants because these two methods prefer to consider STRs with a short sequence context around variants instead of seeking larger tandem repeats with a longer sequence context. TRF and RepeatMasker annotated some variants that were not annotated by other methods because the ‘Simple Repeats’ and ‘RepeatMasker’ tracks of the UCSC Genome Browser contained interrupted STRs, which made STR regions larger and allowed more variations between STR units. When we filtered out interrupted STRs from TRF and RepeatMasker, all these unique TRF and RepeatMasker annotations were removed (S1file: S1 Fig). VarSCAT also made some unique annotations because it not only considered the single reported position of a variant (‘POS’ column in the VCF file) but also considered a range of sequence contexts as variant sites. VarSCAT can also annotate variants located partially in STR regions and variants directly adjacent to STR regions, whereas annotation tools like ANNOVAR only consider the single positions of variants when annotating VCF files with sequence annotations.

**Fig 2.**
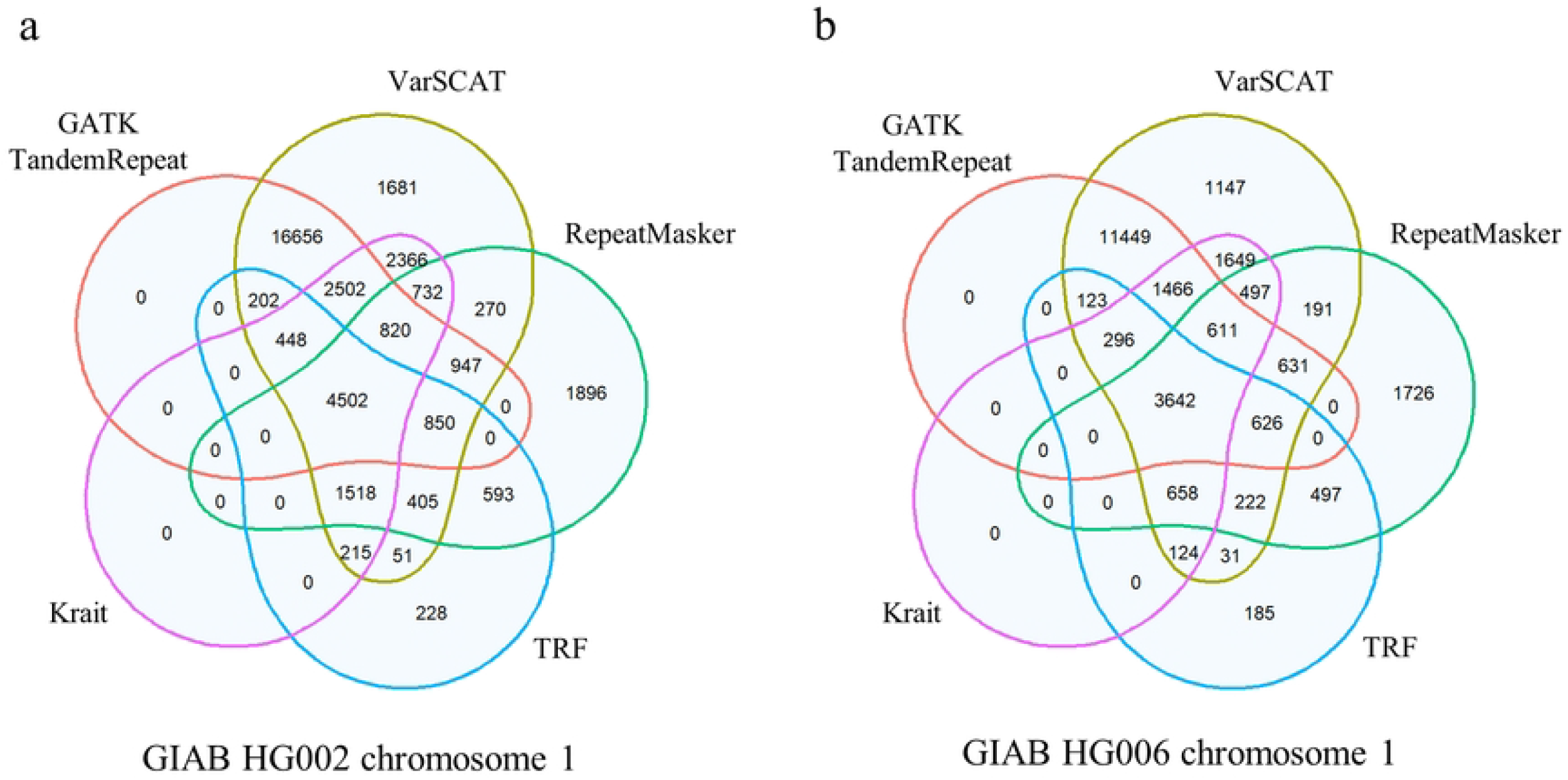
The benchmarking results of the VarSCAT tandem repeat variant annotation module. Benchmarking was performed with small variants in chromosome 1 of **(a)** GIAB HG002, and **(b)** GIAB HG006. GATK ‘TandemRepeat’ function is an annotation method that directly takes a VCF file as the input; Krait is an annotation method for detecting perfect STRs with a reference genome; TRF and RepeatMasker from the UCSC Genome Browser’s ‘Simple Repeats’ and ‘RepeatMasker’ tracks represent a ready-made STR annotation approach. The numbers are counts of variants annotated in STR regions by each tool.

### Ambiguous breakpoint indels and indels in duplicate involve a large number of human indels

To study the proportions of ambiguous breakpoint indels and indels in duplicate in the human genome, we considered indels from the ClinVar database as clinically related indels, and took Platinum Genome NA12878 and NA12877 and GIAB HG002–HG007 as neutral germline indels (Fig 3 and S1 file: Section S4). We defined an indel as an ambiguous breakpoint indel if its 5’- and 3’- aligned positions differed. Our results showed that the majority of indels in the human genome had breakpoint ambiguities (Fig 3A). For the ClinVar database, more than 75% of all indels had breakpoint ambiguities, with 46% and 31% breakpoint-ambiguous deletions and insertions, respectively. For small germline indels in individual human samples, around 90% of all indels had breakpoint ambiguities, with breakpoint-ambiguous deletions and insertions around 45%.

**Fig 3.**
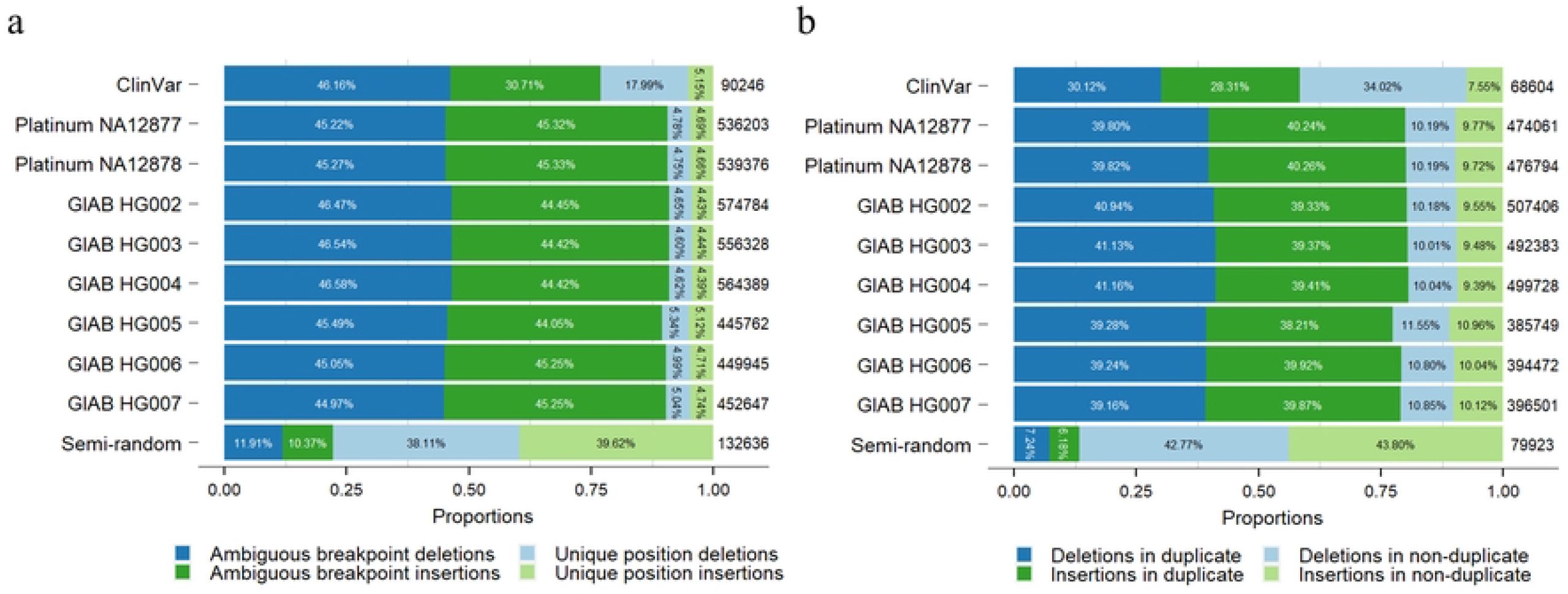
The proportions of ambiguous breakpoint indels and indels in duplicate. The analysis was performed with indels from the ClinVar database, eight high-confidence human individual germline small variant sets, and one semi-random indel set. The proportions of deletions and insertions in different categories are shown separately: **(a)** the proportion of ambiguous breakpoint indels and **(b)** the proportion of indels in duplicate. The numbers on the right of each bar are the numbers of ambiguous breakpoint indels and indels in duplicate for each indel set, respectively.

We further restricted our criteria to analyzing the proportion of indels in duplicate. We defined an indel located in a duplicate if the distance between its 5’- and 3’- aligned positions was larger than the size of the indel; more specifically, if a deletion occurred in a duplication sequence context or higher repeated sequence context (e.g., triplication and quadruplication) and an insertion formed a duplication or higher repeated sequence. Our results showed that most indels in the human genome located in duplicates, that nearly 60% of indels in the ClinVar database and around 80% of indels in the small germline indel sets of eight human individuals (Fig 3B).

Indels in the ClinVar database had lower proportions of both breakpoint-ambiguous indels and indels in duplicate than individual human germline small indel sets. The reason for this is possibly that the average size of indels in the ClinVar database (mean: 56 bp and median: 3 bp) was larger than the average size of human germline small indels (for example, mean: 3 bp and median: 1 bp for Platinum NA12878). The larger indels contained more sequence diversity, thus making the proportions of breakpoint ambiguities and indels in duplicate low.

To further reveal the specificity of the sequence contexts of indels, we created a semi-random small indel set to validate our results (S1 file: Section S4). The total count and size distribution of the semi-random indels were simulated by the indel set of Platinum Genome NA12878, but with indels randomly inserted into the human reference genome GRCh38. The proportions of breakpoint-ambiguous indels and indels in duplicate in the semi-random small indel set showed large differences compared to the real human individual germline small indel sets. Only around 22% and 13% of the semi-random small indels were breakpoint-ambiguous indels and indels in duplicate, respectively. The results were close to the expectations of random selections of four letters, such as for the human genome. Our results demonstrated that human indels appeared in a specific sequence context, and the breakpoint ambiguity issue could not be ignored.

### Characteristics of small variants in the STR regions of the human genome

To study the proportions of variants in STR regions of the human genome in general, we analyzed small variants for 2,548 human individuals from the 1000 Genomes Project that were mapped against GRCh38 (S1 file: Section S5). Our results showed that, on average, 7.1% of individual human germline small variants (including indels) were located in STR regions. For individual human germline small indels, 35.7% of them were located in STR regions (Fig 4). At the superpopulation level, American population, East Asian population, European population, and South Asian population had similar average proportions of small variants and indels in STR regions, while African population had a smaller average proportion but a higher average number of small variants and indels in STR regions compared with the other superpopulations. Similar results were obtained when we analyzed proportions of small variants and indels in STR regions at the subpopulation level (S1 file: S2 Fig).

**Fig 4.**
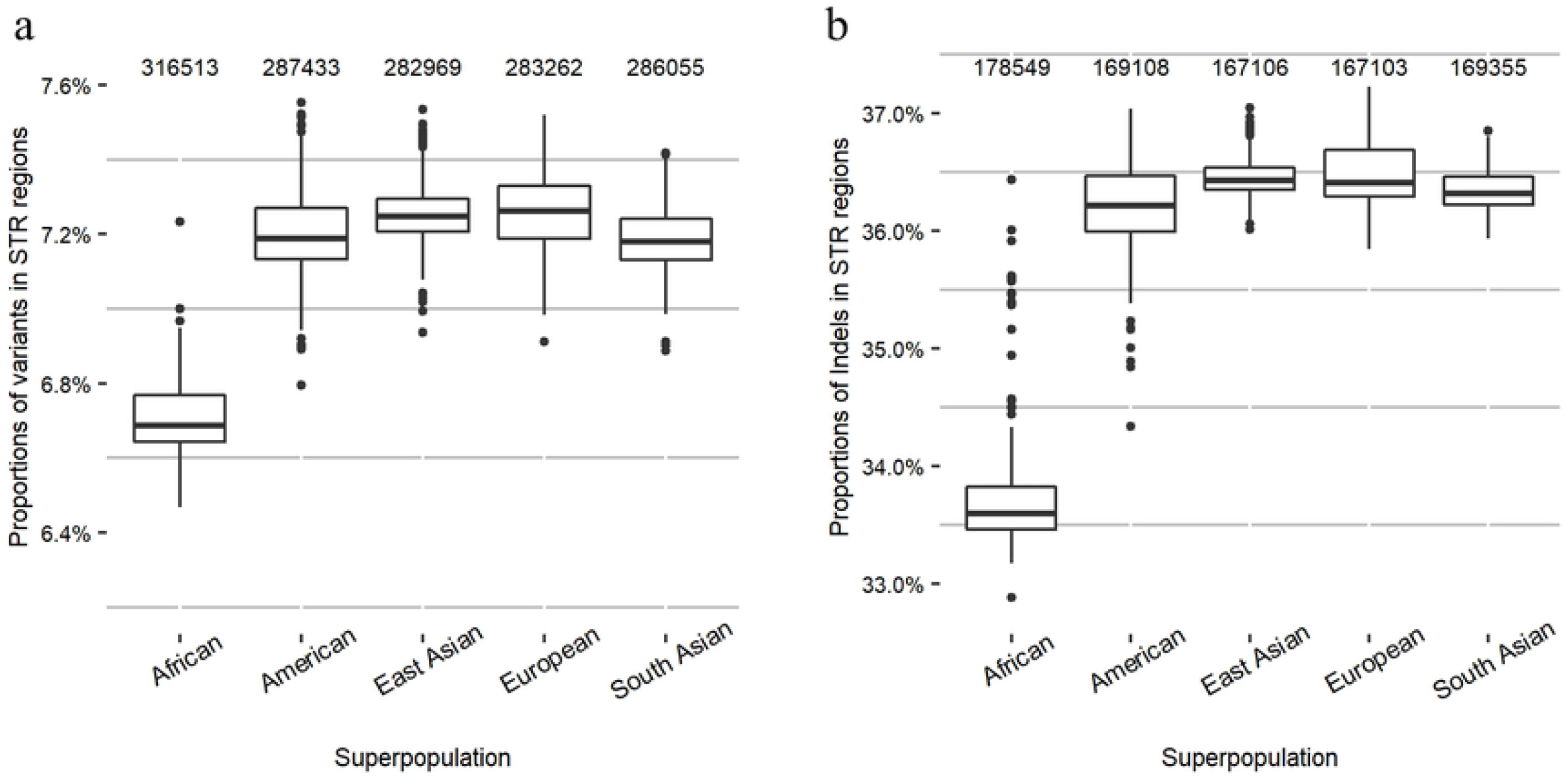
The proportions of small variants and indels in STR regions in different human superpopulations in the 1000 Genomes Project. (a) The proportions of small variants in STR regions and (b) the proportions of small indels in STR regions. The numbers at the top of each boxplot are the average numbers of variants or indels in the STR regions of each superpopulation. African: n = 671; American: n = 348; East Asian: n = 515; European: n = 522; South Asian: n = 492.

To further study the proportions of small variants in STR regions that were shared by the superpopulations, we chose common variants that had at least a 5% minor allele frequency among any of the superpopulations. The results showed that a large proportion (55.2%) of small variants in STR regions were shared by all five superpopulations, and that the proportion was higher than the proportion of small variants in non-STR regions (42.1%; Fig 5). Furthermore, the proportions of small variants in STR regions that were superpopulation specific were smaller (0.8–17.7%) than the proportions of small variants in non-STR regions (1.2–27.5%). This trend was seen in all five superpopulations. The results indicated that small variants in STR regions were more common and more often shared between superpopulations than small variants in non-STR regions.

**Fig 5.**
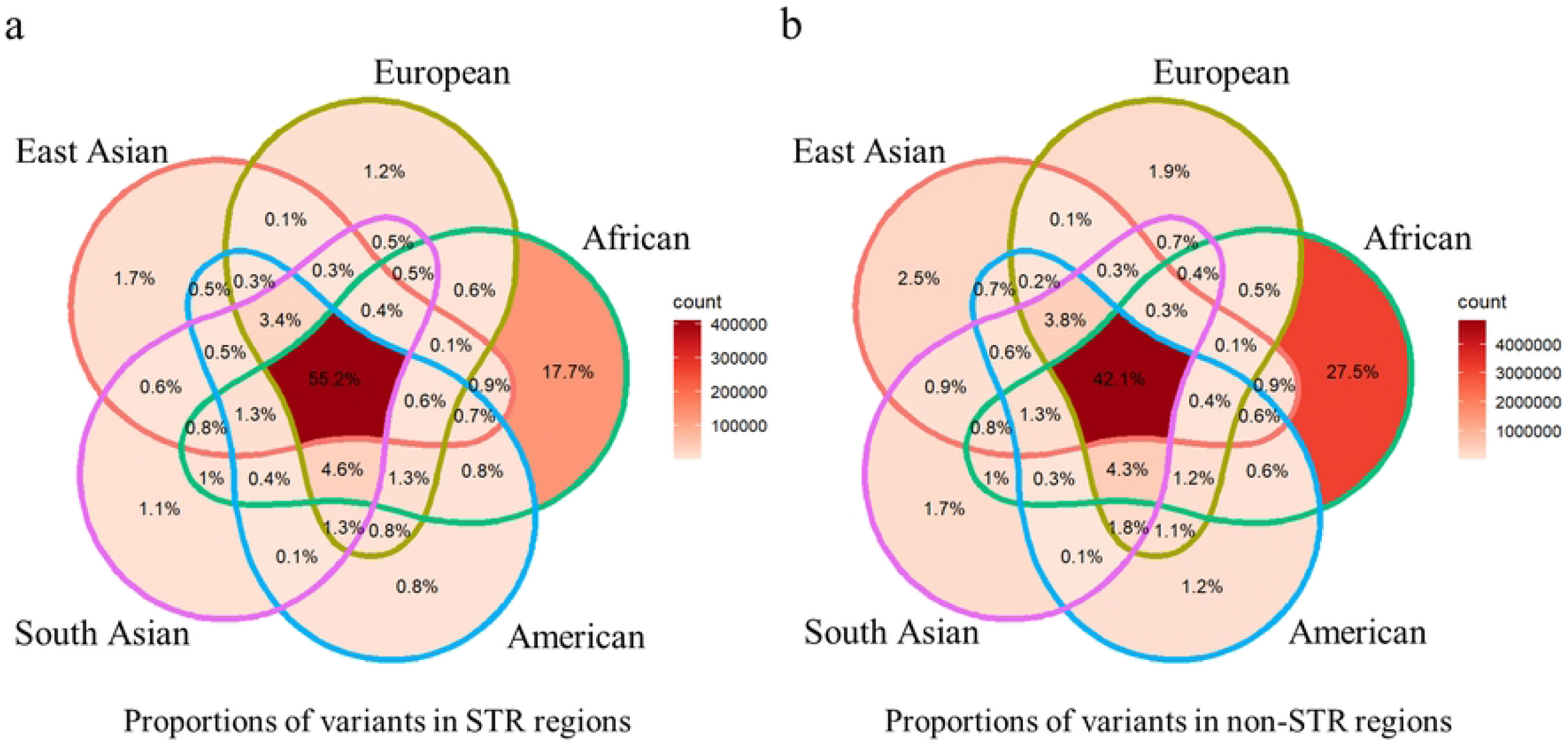
The proportions of small variants in STR and non-STR regions shared by human superpopulations in the 1000 Genomes Project. **(a)** the proportion of small variants in STR regions and **(b)** non-STR regions shared by superpopulations.

To investigate the small variants in the STR regions of single individuals, we analyzed high-confidence small variant sets of two samples from Platinum Genome and six samples from GIAB (Fig 6 and S1 file: Section S5). The results showed that among the small variants in the STR regions, the deletions had the largest proportion (around 45%), followed by insertions of around 40% and single nucleotide variants (SNVs) of around 15%. In contrast, SNVs were the dominant variant type in non-STR regions (above 92%), while both deletions and insertions had low proportions in non-STR regions of around 4%, respectively.

**Fig 6.**
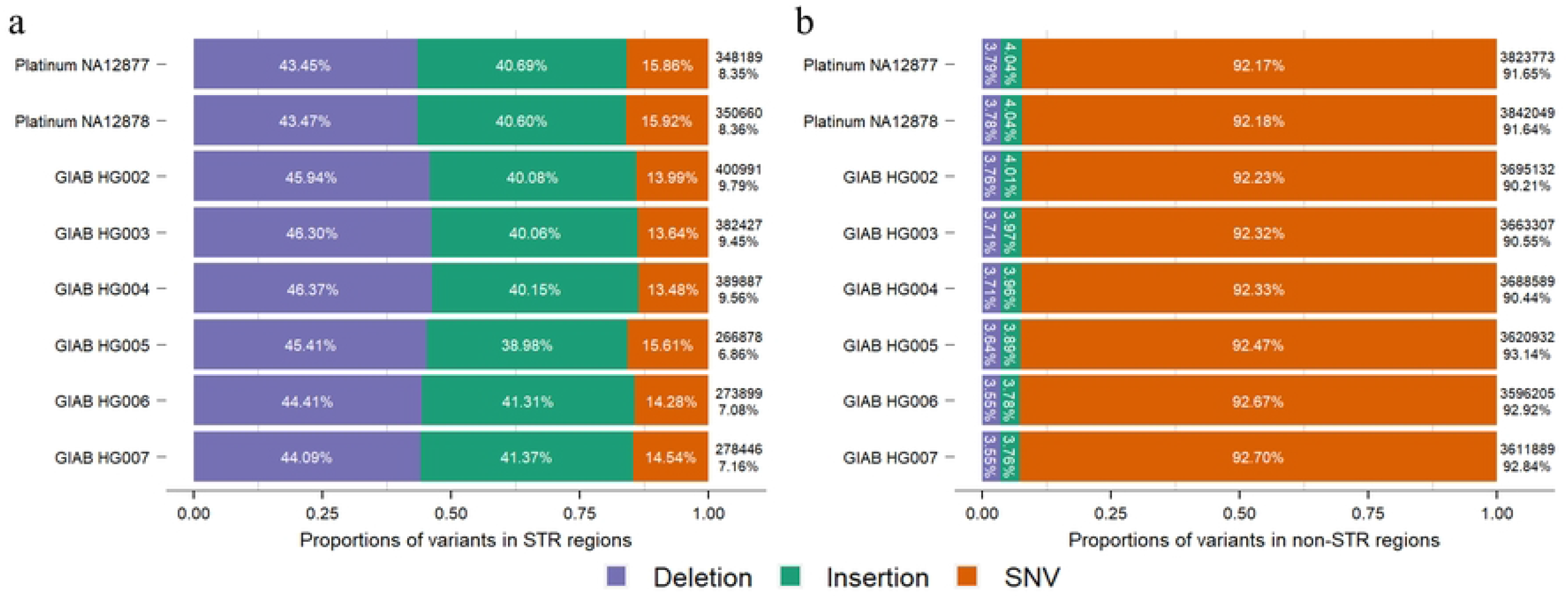
The proportions of different types of small variants in the STR and non-STR regions. **(a)** the proportions of deletions, insertions, and SNVs in STR regions and **(b)** the proportions in non-STR regions. The analysis was performed on high-confidence small variant sets from two individuals from Platinum Genome and six individuals from GIAB. The numbers on the right of each bar are the numbers of STR-region or non-STR-region small variants for the individuals. The percentages on the right of each bar are the proportions of STR-region or non-STR-region small variants among all variants for the individuals.

We further selected small indels in STR regions that spanned only one STR region (on average, 96.7% of total small indels in STR regions in eight individuals) and studied the correlations between indel sizes and STR motif sizes. Our results showed that the majority of small indels in the STR regions were the same size as the STR motif itself (e.g., indels of 2 bp were located predominantly in the dinucleotide STR regions). Indel sizes were also enriched with multiples of STR motif sizes (e.g., dinucleotide STR regions contained indels affecting 2, 4, 6, etc. nucleotides). These findings applied equally to both deletions and insertions (S1 file: S3–S8 Figs). As expected, the counts of STR regions that contained small indels decreased with increases in STR copy numbers and STR motif sizes (S1 file: S9–S14 Figs).

The overall proportion of small variants in the STR regions in eight high-confidence human small variant sets was higher than for the individuals in the 1000 Genomes Project (Figs 4 and 6). This may have been due to improved sequencing techniques and computational variant calling methods that detected high-confidence small variants from difficult regions of the human genome, including STR regions. The 1000 Genomes Project generated variants with the Illumina NGS short-read platform, while GIAB produced high-confidence variant sets with various sequencing platforms, including NGS short-read and third generation, long-read sequencing platforms, as well as variant calling methods based on deep learning approaches. Our results indicated that indels were the most common small variant type in STR regions, and the size distribution of small indels in STR regions had high correlations with the size of the STR motifs.

## Discussion

Sequence contexts can substantially influence genomic variants. Assessing the sequence contexts of variants can help in understanding potential technical issues, such as biologically equivalent indels caused by breakpoint ambiguity, as well as biological significances, such as variants in the STR regions. To learn the sequence contexts of variants using sequencing data, annotation tools such as ANNOVAR can be integrated with other annotation sources to build custom pipelines for annotating variants. However, building such pipelines requires great effort, which can be time-consuming and difficult. Hence, we developed a single command-line-based computational tool VarSCAT, which takes a VCF file and a reference sequence FASTA file as inputs. VarSCAT has various functions for annotating breakpoint ambiguities, flanking bases of variants, HGVS nomenclature, distances between adjacent variants, and tandem repeat regions of variants, as well as for extracting reference DNA and alternative DNA sequences by considering all the variants within a user-defined region. The VarSCAT results provide users with valuable information about the sequence contexts of variants, which can be used for such purposes as variant calling result filtration and variant nomenclature.

To demonstrate the utility of VarSCAT, we used its flexibility to analyze high-confidence human germline small variant sets from Platinum Genome, GIAB, and the 1000 Genomes Project, and clinically related indels from the ClinVar database to study the sequence contexts of variants. Our benchmarking results for the annotation of variants in STR regions showed substantial discordance between the different methods and strategies, indicating that the current methods and strategies may underestimate the proportions of variants in STR regions. Some STRs not be annotated by other methods in our benchmarking may be because the strict criteria that filter them out or because they are part of large tandem repeats and are thus ignored. Our results showed that the majority of human indels have breakpoint ambiguities, consistent with a previous study [23].

Although STRs occupy only about 3% of the human genome [64], our study showed that on average, more than 7% of human germline small variants were in STR regions—especially small indels, which were the most dominant variant type in STR regions. Small variants in STR regions were more common than in non-STR regions across human superpopulations, and the sizes of small indels in STR regions correlated highly with the sizes of the STR motifs. Our results agreed with previous research indicating that DNA strand slippage is the main molecular mechanism underpinning indel generation [65], and that the current African populations have larger genetic variations than the non-African populations [13].

Although VarSCAT can annotate variant sequence contexts comprehensively in a high-throughput manner, our study and results still have some limitations. Due to the inconsistent criteria for interrupted STRs, such as mismatches and indels tolerance, we limited our research to perfect STRs, which may have led to an underestimation of the proportions of variants in STR regions. STR regions annotated by VarSCAT may not always have biological variant significance, and further validations are needed, such as studies of the functions of STRs and variants. The current variant annotation methods that take VCF files as inputs only consider the initial reported positions of variants without considering variants, especially indels as effected regions on genome, thus, they may miss annotations if there is only a partial overlap between variants and STR regions. Although VarSCAT can annotate variants that only partially overlap with an STR, we focused on high-confidence individual human small variant sets for analysis. The high-confidence small variant sets had high precision and covered much of the human genome, but they still excluded some difficult regions that were enriched with STRs and limited the sizes of variants due to difficulties in variant genotyping [47–49].

The data we used from the 1000 Genomes Project were the most recent variant call sets, based on the current human reference genome assembly GRCh38. The data contained 2,548 samples, which gave us an adequate sample size to study the general proportions of small variants in the STR regions of human individuals. However, the data contained only biallelic variants. Although the majority of variants in Phase Three of the 1000 Genomes Project are biallelic [51], we still may have systematically underestimated the proportions of small variants in STR regions because these variants are often highly multiallelic [66]. With improved sequencing techniques, such as long-read sequencing and corresponding variant calling methods, the STR-enriched regions, such as centromeres and telomeres, could be precisely sequenced, and the variations among individuals could be genotyped accordingly, which has the potential to further improve studies of variants in STR regions using VarSCAT.

## Methods and Materials

### VarSCAT annotation modules

VarSCAT is a computational tool which takes a VCF file and a reference sequence as inputs to annotate the sequence contexts of genomic variants. First all the input variants are processed by variant normalization to splits any multiallelic variants into biallelic variants and then normalizes them. The ambiguous variant annotation module annotates variants related with breakpoint ambiguities and the tandem repeat region variant annotation module can annotate putative tandem repeat regions that contain variants. These two modules and their functions can be used together or separately. The design of each module is described as follows.

### Variant normalization

For VarSCAT’s first analytical step, we built a module that normalizes input variants parsimonious and left aligned after separating any potential multiallelic variants into biallelic variants. A variant is parsimonious if (and only if) it is represented in as few nucleotides as possible without an allele of zero length [41]. A variant is left aligned if (and only if) it is no longer possible to shift its position to the left while keeping the length of all its alleles constant [41]. This variant normalization module can be used to analyze input variants in a standardized way, without further concerns about multiple representations of a variant.

VarSCAT requires a VCF input file. The variant information for chromosomes (‘CHROM’ column), positions (‘POS’ column), reference alleles (‘REF’ column), alternative alleles (‘ALT’ column), and genotypes (‘GT’ section in the ‘INFO’ column) are read by the PyVCF package [42]. The variant normalization algorithm is similar to the vt tool [41]. For normalization, VarSCAT extracts indels and multi-nucleotide variants (MNVs) with REF or ALT > 1 bp. SNVs have single positions with single nucleotide substitutes and do not need to be normalized. First, for indels and MNVs, if the rightmost sequence nucleotides of REF and ALT are the same, VarSCAT trims this common last nucleotide until the uncommon nucleotides are met. During this trimming, if the sequence length of any one of REF or ALT is zero, then both REF and ALT are extended by one nucleotide to the left based on the reference genome sequence, and the position of the variant is updated. After trimming to the rightmost sequence nucleotides, VarSCAT examines the leftmost nucleotides of the variants. If both REF and ALT lengths are longer than one and have common nucleotides, then the tool trims the leftmost nucleotides of REF and ALT and updates the position of the variant. After the normalization process, the chromosomes, positions, REFs, ALTs, and genotypes of all the input variants are stored.

### Ambiguous variant annotation module

With the ambiguous variant annotation module, VarSCAT returns each variant record as a row in a text file. Each record from this module contains basic variant information (chromosome, position, REF, ALT, and genotype) and optional information, including 5’-aligned position, 3’-aligned position, 3’ edge position (the rightmost base position of a variant), the flanking bases of REF and ALT, the sequence variant nomenclature recommended by HGVS (version: 20.05) [43], and the distance to the next variant on 3’ direction by activating additional parameters. Furthermore, if the user provides a genomic region, VarSCAT can output the region-specific reference DNA sequence, the mutated DNA sequence, and its reverse complement sequence in an FASTA file.

Because a point nucleotide change occurs only at a fixed position, there is no breakpoint ambiguity for SNV, MNV, and complex indel (for example, REF: CATTC, ALT: G). For the variants that contain a point nucleotide change, the 5’-aligned and 3’-aligned positions are the positions of the first nucleotide of the variant (the same as the position of the variant), and the 3’ edge position is the position of the last nucleotide of the variant. For indels, we first compute the 3’ edge positions. VarSCAT first extracts the indel pattern, which is the leftmost nucleotide trimmed sequence of REF or ALT for deletion and insertion, respectively. If the pattern of an indel with length *n* is a nucleotide subsequence (*a*_*1*_, *a*_*2*_, *a*_*3*,_ …, *a*_*n*_), the adjacent reference sequence in the 3’ direction to the indel is a nucleotide sequence (*r*_*1*_, *r*_*2*_, *r*_*3*,_ …) with genomic coordinates (*c*_*1*_, *c*_*2*_, *c*_*3*,_ …), so the initial 3’ edge position of the indel is the genomic coordinate *c*_*1*_-1. VarSCAT checks whether the leftmost nucleotide of the indel pattern *a*_*1*_ is the same nucleotide as the first 3’-direction reference sequence nucleotide *r*_*1*_. If *a*_*1*_ is the same nucleotide as *r*_*1*_, then the indel pattern permutes into (*a*_*2*_, *a*_*3*_…*a*_*n*_, *a*_*1*_) and the 3’ edge position updates to the genomic coordinate *c*_*2*_-1. In the next round, the updated leftmost nucleotide of the indel pattern *a*_*2*_ is compared with the second 3’ direction reference sequence nucleotide *r*_*2*_. If they are the same, VarSCAT continues the variant pattern permutation, and the 3’ edge position is updated; if they are not the same, VarSCAT terminates the process and reports the 3’ edge position. For deletion, the 5’-aligned position is the position of the leftmost possible nucleotide of the deleted sequence on the reference coordinate, and the 3’-aligned position is the position of the rightmost possible nucleotide of the deleted sequence on the reference coordinate (3’ edge position) minus the length of the deletion pattern; for insertion, the 5’-aligned position is the position of the leftmost possible 5’ direction adjacent nucleotide of the inserted sequence on the reference coordinate (the same as the position of variant), the 3’-aligned and 3’ edge positions are the same as the position of the rightmost possible 5’ direction adjacent nucleotide of the inserted sequence on reference coordinate. With the 5’-aligned and 3’ edge positions, the 3’ direction distance to the next variant from the input VCF can be calculated.

We computed the HGVS nomenclature of variants followed by the HGVS recommendations (detailed in https://varnomen.hgvs.org/recommendations/DNA/) with a custom algorithm and script using the 5’-aligned and 3’-aligned positions. Also, if the ambiguity region of a variant is known, VarSCAT can output the variant flanking bases, including the potential ambiguity sequences of REF and ALT. If the user provides a genomic region, VarSCAT can extract a reference DNA sequence from this region together with variants within this region to create the mutated DNA sequence. The reverse complement sequence of a mutated DNA sequence was created using the Biopython package [44].

### Tandem repeat region variant annotation module

VarSCAT uses a local sequence context comparison algorithm designed according to the idea of tandem repeat definition to explore the tandem repeat regions that either overlap a variant or are adjacent to a variant.

#### Initiation of a variant site

Let *min_unit* be the minimum size of a conserved repeat motif, and *max_unit* be the maximum size of a conserved repeat motif, defined by the user or by default. For each input variant that does not contain ‘N’, VarSCAT first defines the variant site. For deletion, the variant site is the coordinates of the deleted sequences relative to the reference sequence; for insertion, the variant site is the coordinates of two nucleotides on the reference sequence, located in the 5’ and 3’ directions adjacent to the inserted sequence, respectively; for SNV and MNV, the variant site is the coordinates of the substituted sequence relative to the reference sequence. For a known variant site, VarSCAT creates a window size set *W* ={*w*_*1*_, *w*_*2*_, *w*_*3*,_ …} in which *w*_*i*_ is a certain window size and *W* contains window sizes with a range of *min_unit* ≤ *w*_*i*_ ≤ *max_unit*. For each certain window size *w*_*i*_, VarSCAT creates a set of nucleotide subsequences *R* ={*r*_*1*_, *r*_*2*_, *r*_*3*,_ …}, in which *r*_*i*_ is subsequence with size *w*_*i*_ as a candidate conserved repeat motif and *R* contains all the possible subsequences with at least one base pair overlapped or directly adjacent to the variant site in the 5’ to 3’ direction.

#### The local sequence context comparison algorithm for repeat units searching

If *r*_*i*_ with size *w*_*i*_ is one candidate conserved repeat motif. VarSCAT first takes the 5’ direction to search for potential repeat units. The algorithm creates a set of nucleotide subsequences *S* ={*s*_*0*_, *s*_*1*_, *s*_*2*,_ …, *s*_*wi-1*_}. Nucleotide subsequences in *S* have the same size of *w*_*i*_ and each *s*_*i*_ is a nucleotide subsequence created by moving window *w*_*i*_ 1 bp away from *r*_*i*_ in the 5’ direction per moving step. Each moving step can form a gap, and the gap distance between *r*_*i*_ and *s*_*i*_ is from 0 bp to a maximum of (*w*_*i*_–1) bp or a user-defined fixed maximum gap tolerated distance. Each *s*_*i*_ is treated as a potential repeat unit and Set *S* ensures that VarSCAT captures all subsequences as potential repeat units within the maximum gap-tolerated distance. With all possible *s*_*i*_ in *S*, a similarity comparison between *r*_*i*_ and *r*_*i*_ is performed. Due to the same string lengths between *r*_*i*_ and *s*_*i*_, the number of common bases at each string location is used as the similarity comparison score. For all possible *s*_*i*_ and their corresponding similarity comparison scores, the *s*_*i*_ with the highest score is chosen, and a corresponding gap distance is determined (if there are equal best scores, VarSCAT will choose the *s*_*i*_ with the smallest gap distance). If *s*_*i*_ with the highest similarity comparison score passes the default or user-defined similarity threshold with *r*_*i*_, *s*_*i*_ is treated as one repeat unit, and the gap distance and the number of mismatched nucleotides between *r*_*i*_ and *s*_*i*_ are stored before the next round is executed. For the next round, a new set of subsequences *S* is created in the 5’ direction of *s*_*i*_ and the same local sequence context comparison and criteria are applied to search the next repeat unit *s*_*i + 1*_. The process is terminated when all the possible elements in *S* in the *n*th round fail to pass the similarity threshold with *r*_*i*_, and the same process is performed but in the 3’ direction from *r*_*i*._ After searching and collecting all possible repeat units in both the 5’ and 3’ directions, a candidate tandem repeat region of motif *r*_*i*_ is formed, and the numbers of matched nucleotides, mismatched nucleotides, gaps, and the copy number in this tandem repeat region are stored.

The VarSCAT results had similar underpinning technical concepts, such as ‘Match’, ‘Mismatch’, and ‘Gap’ to a global pairwise alignment based on the Needleman–Wunsch algorithm [45]. The VarSCAT results can be seen in terms of a Needleman–Wunsch algorithm that aligns the VarSCAT-defined tandem repeat region of motif *r*_*i*_ with an ideal perfect tandem repeat region of the same motif *r*_*i*_ under the same copy number. If the copy number of the candidate tandem repeat region is larger than the default or user-defined minimum copy number, the alignment score (Equation. 1) is calculated for the candidate tandem repeat region with the default or user defined match score (MS), mismatch score (MIS), and gap score (GS). We defined a repeat score as the alignment score divided by the length of the candidate tandem repeat region and then multiplied by the corresponding copy number (Equation. 2). The conserved repeat motif *r*_*i*_, copy number, size of the conserved repeat pattern, start and end positions of the tandem repeat region, repeat score, alignment score, match percentage, mismatch percentage, gap percentage, and history records from every local sequence context comparison round for the start and end positions, repeat patterns, number of mismatches, and gaps of all the candidate tandem repeat regions are stored for post-processing quality control. Due to DNA having four different types of bases (A, T, G and C), during the first four rounds, there may be no consensus nucleotide pattern as a potential motif for local sequence context comparison. Therefore, during the first four rounds, *r*_*i*_ is the subsequence that created for the variant site. After the fifth round, *r*_*i*_ is updated to the consensus sequence motif (*r*_*i*_, *s*_*i*_, *s*_*i2*_, *s*_*i3*,_ …, *s*_*in*_). Every round after the fifth round uses the updating consensus sequence motif for the local sequence context comparison, and the alignment results of the first four rounds are replaced with the conserved sequence motif.

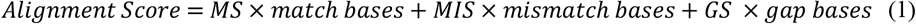

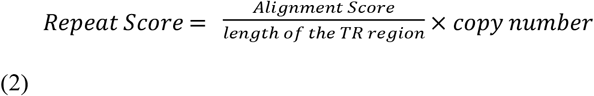

#### Post-processing quality control

Post-processing quality control is applied using the default or user-defined parameters of 1) the minimum alignment score to control the minimum size of a tandem repeat region and 2) the minimum match percentage to control the repeat purity of a tandem repeat region. A tandem repeat region for a variant is stored if the tandem repeat region passes the quality threshold. If a variant has at least one candidate tandem repeat region after searching but no tandem repeat passes the quality threshold, a trimming algorithm is applied to the tandem repeat region with the best alignment score. We assumed that the 5’ and 3’ tails of a tandem repeat region would share less similarity and contain more gaps between the conserved repeat pattern, which could cause the alignment score and the match percentage to decrease below the quality threshold. VarSCAT trims the historical records of the repeat units by searching the results for both the 5’ and 3’ tails of a candidate tandem repeat region until the copy number is less than the threshold. If, during the trimming process, the match percentage and alignment score exceed the quality threshold, the trimming process is terminated, and the corresponding values of the candidate tandem repeat region are recalculated. Any candidate tandem repeat region of a variant site that meets the criteria is stored with its conserved repeat motif, copy number, repeat score, percentage of matches, percentage of mismatches, percentage of gaps, and repeat region for further analysis.

#### Remove redundancy

After all potential candidate tandem repeat regions of a variant site are discovered, it is still possible for redundant tandem repeat regions to be recorded. The conserved repeat motifs of a redundant repeat region are usually multiples of each other, but only the most concise conserved repeat motif is needed. VarSCAT first ranks all the candidate tandem repeat regions with repeat scores in descending order, favoring the tandem repeat with the highest copy number. Starting with the highest repeat score candidate tandem repeat, all candidate tandem repeats are compared in a pairwise manner. We defined a redundancy ratio to filter out redundant tandem repeat regions. The redundancy ratio is the ratio of the size of overlapped regions and the maximum covered regions of the two pairwise compared tandem repeats on reference sequence. If the redundancy ratio is > 0.5, meaning that the size of an overlapped region is larger than half of the maximum covered region of two tandem repeats, VarSCAT considers one of the two tandem repeats as a redundant tandem repeat. For each tandem repeat in pairwise comparison, VarSCAT calculates the nucleotide frequency difference between its conserved repeat motif and the whole tandem repeat region. The tandem repeat with the smallest nucleotide frequency difference remains, and the other tandem repeat is dropped. For all remaining tandem repeats, the GC% is calculated and they are reported as the final tandem repeats for the variant.

### Data sets

We investigated several high-confidence genomic variant data sets that either included variants with clinical significance or represented the natural distribution of human germline variants. We selected variants from the ClinVar database [46], two high-confidence small variant calls for human individuals NA12877 and NA12878 from Platinum Genomes [47], six high-confidence small variant calls (v4.2.1) for human individuals HG002-HG007 from the GIAB Consortium [48,49], and the integrated phased biallelic variant sets on GRCh38 for 2,548 human individuals from the 1000 Genomes Project [50,51]. An extended description of the data selection and processing is provided in S1 file: Sections S3–S5.

#### ClinVar database

The ClinVar database (data archiving date: 2022/01/09) is a public archive hosted by the National Center for Biotechnology Information, which holds the relationships among human genome variations and phenotypes with supporting clinical evidence. The records in this database are alleles that have been mapped to reference sequences and reported according to the HGVS nomenclature standard, together with their clinical significance or supporting evidence about the effects of the variations [46]. The indels on chromosomes 1–22, sex chromosomes, and the mitochondrial chromosome in the ClinVar database were selected for the indel analysis of breakpoint ambiguity and duplicate.

#### Platinum Genome

Platinum Genome is a high-confidence human small variant set that contains two human individuals (NA12877 and NA12878) from the CEPH pedigree 1463. The high-confidence variant set was generated using PCR-free whole genome sequencing for four grandparents, two parents (NA12877 and NA12878), and 11 children, and the calling variants in each genome using several publicly available, highly accurate variant calling algorithms. Platinum Genome used haplotype transmission information for this pedigree to create a high-confidence variant set containing 4.7 million SNVs and 0.7 million small indels that ranged from 1–50 bp [47]. **GIAB**: The GIAB high-confidence small variant sets contain seven human samples (NA12878 and two son/father/mother trios of Ashkenazi Jewish and Han Chinese ancestry). High-confidence variant sets were generated using an integration pipeline of computational tools with sequencing data from multiple technology platforms, including both short-read and long-read sequencing platforms. In our study, we used GIAB high-confidence variant sets v4.2.1 of the trios of Ashkenazi Jewish and Han Chinese ancestry (HG002–HG007), in which more SNVs and indels located in difficult-to-map regions were added and covered 92% of the autosomal GRCh38 assembly [48,49]. **The 1000 Genomes Project**: The 1000 Genomes Project is a comprehensive catalogue of common human genetic variations with allele frequencies of at least 1% in the populations studied. The data were produced by applying whole-genome sequencing for the human reference genome GRCh38 [50] to a diverse set of individuals from multiple populations. In our study, we used a set of biallelic SNVs and indels from 2,548 samples of 26 populations that were generated by the 1000 Genomes Project based on the human reference genome GRCh38. The variant call set was produced using a multi-caller approach, which integrated the call sets before final genotyping and phasing [51].

### Benchmarking of STR region variant annotation

Variants in VCF files for chromosome 1 of two human individuals (HG002 and HG006) from GIAB were selected for benchmarking the variants annotation of STR regions after splitting multiallelic variants (GATK v4.1.9.0, ‘--split-multi-allelics’) and left alignment (vt v0.57712, ‘left-aligning’). The ‘Simple Repeat’ and ‘RepeatMasker’ tracks of chromosome 1 of human reference hg38, representing TRF and RepeatMasker, respectively, were downloaded from the UCSC genome browser (download date 11 February 2022). The STR records generated by Krait (v1.3.3) were limited to perfect STRs. All these STR records were then filtered based on the minimum lengths and copy numbers of STRs, as mentioned in the ‘Results’ section. We used a custom R script to create BED files for these STR records and used ANNOVAR (version: ‘$Date: 2019-10-24’) to annotate variants in VCF format. The VarSCAT and GATK TandemRepeat (v4.1.9.0) annotation results, which both directly took VCF files as inputs, were limited to perfect STRs according to our criteria of the minimum length and copy numbers, as mentioned in the ‘Results’ section (details are given in S1 file: Section S2).

### Data availability statement

VarSCAT is an open-source tool and released under the GPLv3 license. The source code of VarSCAT and documentation is available at https://github.com/elolab/VarSCAT. The codes of analysis of this manuscript are available at https://github.com/elolab/VarSCAT-analysis. In addition, the source code of VarSCAT and the codes of data analysis of this manuscript can be accessed in Zenodo at https://doi.org/10.5281/zenodo.7018364 [67]. The variant set of ClinVar database analyzed during the current study are available at https://www.ncbi.nlm.nih.gov/clinvar/. The high-confidence variant sets of Platinum Genome analyzed during the current study are available at https://www.illumina.com/platinumgenomes.html. The high-confidence variant sets from GIAB analyzed during the current study are are available at https://www.nist.gov/programs-projects/genome-bottle. The 2,548 human individual variant sets from the 1000 Genome Project analyzed during the current study are available at https://www.internationalgenome.org/.

## Funding

**N**ing Wang received funding from Turku University Foundation, and University of Turku Graduate School (UTUGS). Prof. Elo reports grants from the European Research Council ERC (677943), European Union’s Horizon 2020 research and innovation programme (955321), Academy of Finland (310561, 314443, 329278, 335434, 335611 and 341342), and Sigrid Juselius Foundation during the conduct of the study. Our research is also supported by Biocenter Finland, and ELIXIR Finland.

## Acknowledgements

We thank Salla Rusanen for her valuable comments and António Sousa for testing VarSCAT.

## Competing interests

The authors declare that they have no competing interests.

## Author Contributions

NW and SK originated the study. NW designed the research, developed the method, implemented the software, performed benchmarking analysis, analyzed high-confidence variant sets, performed visualizations, and drafted the manuscript. NW, SK and LLE discussed the results. SK and LLE supervised the work. All authors contributed to manuscript writing. All authors read and approved the manuscript.

## Corresponding authors

Correspondence to Ning Wang and Laura L. Elo.

## Supporting information

**S1 Fig. The benchmarking results of the VarSCAT tandem repeat variant annotation module with only perfect STRs from TRF and RepeatMasker**. The benchmarking was performed with small variants of chromosome 1 of **(a)** GIAB HG002, and **(b)** GIAB HG006. The numbers are the counts of variants annotated by each tool.

**S2 Fig. The proportions of small variants and indels in STR regions in different human subpopulations of the 1000 Genomes Project. (a)** The proportions of small variants in STR regions and **(b)** the proportions of small indels in STR regions. The numbers at the top of each boxplot are the average numbers of variants in the STR regions for each subpopulation. ACB, African Caribbean in Barbados, GWD, Gambian in Western Division Mandinka, ESN, Esan in Nigeria, MSL, Mende in Sierra Leone, YRI, Yoruba in Ibadan Nigeria, LWK, Luhya in Webuye Kenya, ASW, People with African Ancestry in Southwest USA, PUR, Puerto Ricans in Puerto Rico, CLM, Colombians in Medellin Colombia, PEL, Peruvians in Lima Peru, MXL, People with Mexican Ancestry in Los Angeles CA USA, CHS, Southern Han Chinese, CDX, Chinese Dai in Xishuangbanna China, KHV, Kinh in Ho Chi Minh City Vietnam, CHB, Han Chinese in Beijing, China, JPT, Japanese in Tokyo Japan, GBR, British in England and Scotland, FIN, Finnish in Finland, IBS, Iberian Populations in Spain, CEU, Utah residents with Northern and Western European ancestry, TSI, Tuscans in Italy, PJL, Punjabis in Lahore Pakistan, BEB, Bengalis in Bangladesh, STU, Sri Lankan Tamils in the UK, ITU, Indian Telugu in the UK, GIH, Gujarati Indians in Houston TX USA.

**S3 Fig. Distributions of the sizes of small indels in STR regions with an STR motif size of 1 bp**.

**(a)** Platinum NA12877, **(b)** Platinum NA12878, **(c)** GIAB HG002, **(d)** GIAB HG003, **(e)** GIAB HG004, **(f)** GIAB HG005, **(g)** GIAB HG006, and **(h)** GIAB HG007. Deletions and insertions are shown in blue and green, respectively.

**S4 Fig. Distributions of the sizes of small indels in STR regions with an STR motif size of 2 bp**.

**(a)** Platinum NA12877, **(b)** Platinum NA12878, **(c)** GIAB HG002, **(d)** GIAB HG003, **(e)** GIAB HG004, **(f)** GIAB HG005, **(g)** GIAB HG006, and **(h)** GIAB HG007. Deletions and insertions are shown in blue and green, respectively.

**S5 Fig. The distributions of the sizes of small indels in STR regions with STR motif size 3bp**.

**(a)** Platinum NA12877, **(b)** Platinum NA12878, **(c)** GIAB HG002, **(d)** GIAB HG003, **(e)** GIAB HG004, **(f)** GIAB HG005, **(g)** GIAB HG006, **(h)** GIAB HG007. Deletions and insertions are shown as blue and green, respectively.

**S6 Fig. Distributions of the sizes of small indels in STR regions with an STR motif of size 4 bp**.

**(a)** Platinum NA12877, **(b)** Platinum NA12878, **(c)** GIAB HG002, **(d)** GIAB HG003, **(e)** GIAB HG004, **(f)** GIAB HG005, **(g)** GIAB HG006, and **(h)** GIAB HG007. Deletions and insertions are shown in blue and green, respectively.

**S7 Fig. Distributions of the sizes of small indels in STR regions with an STR motif size of 5 bp**.

**(a)** Platinum NA12877, **(b)** Platinum NA12878, **(c)** GIAB HG002, **(d)** GIAB HG003, **(e)** GIAB HG004, **(f)** GIAB HG005, **(g)** GIAB HG006, and **(h)** GIAB HG007. Deletions and insertions are shown in blue and green, respectively.

**S8 Fig. Distributions of the sizes of small indels in STR regions with an STR motif size of 6 bp**.

**(a)** Platinum NA12877, **(b)** Platinum NA12878, **(c)** GIAB HG002, **(d)** GIAB HG003, **(e)** GIAB HG004, **(f)** GIAB HG005, **(g)** GIAB HG006, and **(h)** GIAB HG007. Deletions and insertions are shown in blue and green, respectively.

**S9 Fig. Distributions of copy numbers of motif-size 1 bp STRs containing variants. (a)** Platinum NA12877, **(b)** Platinum NA12878, **(c)** GIAB HG002, **(d)** GIAB HG003, **(e)** GIAB HG004, **(f)** GIAB HG005, **(g)** GIAB HG006, and **(h)** GIAB HG007.

**S10 Fig. Distributions of copy numbers of motif-size 2 bp STRs containing variants. (a)** Platinum NA12877, **(b)** Platinum NA12878, **(c)** GIAB HG002, **(d)** GIAB HG003, **(e)** GIAB HG004, **(f)** GIAB HG005, **(g)** GIAB HG006, and **(h)** GIAB HG007.

**S11 Fig. Distributions of copy numbers of motif-size 3 bp STRs containing variants. (a)** Platinum NA12877, **(b)** Platinum NA12878, **(c)** GIAB HG002, **(d)** GIAB HG003, **(e)** GIAB HG004, **(f)** GIAB HG005, **(g)** GIAB HG006, and **(h)** GIAB HG007.

**S12 Fig. Distributions of copy numbers of motif-size 4 bp STRs containing variants. (a)** Platinum NA12877, **(b)** Platinum NA12878, **(c)** GIAB HG002, **(d)** GIAB HG003, **(e)** GIAB HG004, **(f)** GIAB HG005, **(g)** GIAB HG006, and **(h)** GIAB HG007.

**S13 Fig. Distributions of copy numbers of motif-size 5 bp STRs containing variants. (a)** Platinum NA12877, **(b)** Platinum NA12878, **(c)** GIAB HG002, **(d)** GIAB HG003, **(e)** GIAB HG004, **(f)** GIAB HG005, **(g)** GIAB HG006, and **(h)** GIAB HG007.

**S14 Fig. Distributions of copy numbers of motif-size 6 bp STRs containing variants. (a)** Platinum NA12877, **(b)** Platinum NA12878, **(c)** GIAB HG002, **(d)** GIAB HG003, **(e)** GIAB HG004, **(f)** GIAB HG005, **(g)** GIAB HG006, and **(h)** GIAB HG007.

**S1 File. Supplementary methods descriptions**.

## Notes

### Competing Interest Statement

The authors have declared no competing interest.

